# Affiliative social relationships and coccidian oocyst excretion in a cooperatively breeding bird species

**DOI:** 10.1101/159129

**Authors:** Claudia A.F. Wascher, Daniela Canestrari, Vittorio Baglione

## Abstract

In group living animals, behavioural interactions with conspecifics strongly modulate an individual’s physiological stress response. Stable social relationships may reduce an individual’s stress response, which in turn can affect the immune system and health. Ultimately, positive health effects of stable social bonds may contribute to maintain group living. We investigated whether, in cooperatively breeding carrion crows (*Corvus corone*), the quality of social relationships correlates with coccidian oocyst and nematode eggs excretion. We repeatably collected behavioural data on dyadic social interactions and individual droppings to quantify parasite eggs and oocysts from 36 individuals in a captive population of carrion crows in northern Spain. Individuals with strong social bonds, living with more relatives and in larger groups excreted a significantly smaller proportion of droppings containing coccidian oocysts. The probability to excrete droppings containing nematode eggs was not affected by social factors. The relationship between social interactions and coccidian oocyst excretion is consistent with the idea that high quality social relationships can positively affects individual’s health, setting the stage for the evolution of stable social living.

## Introduction

In group living animals the social environment directly affects an individual’s physiology and health (DeVries, Glasper, & Detillion, 2003; Hanley & Stamps, 2002; Holst, 1998; Sheridan, Dobbs, Brown, & Zwilling, 1994). Adverse effects are caused, for example, by increased competition and aggressive behaviour (Chester, Bonu, & Demas, 2010; Hawley, Lindström, & Wikelski, 2006), which affect especially low-ranking individuals (Ungerfeld & Correa, 2007; Zuk, Kim, Robinson, & Johnsen, 1998). Further, large group sizes increase the exposure to directly transmitted parasites (Côté & Poulin, 1995; Loehle, 1995) and social interaction patterns can also facilitate pathogen transmission (Drewe, 2010; Fouchet et al., 2012).

Affiliative behaviour and reliable social allies, on the other hand, have a stress-reducing effect (Frigerio, Weiss, Dittami, & Kotrschal, 2003; Sachser, Dürschlag, & Hirzel, 1998; Stöwe et al., 2008; Young, Majolo, Schülke, & Ostner, 2014). In several mammalian species, stable social bonds have a fitness enhancing effect (Schülke, Bhagavatula, Vigilant, & Ostner, 2010; Schülke et al., 2010; Silk et al., 2009, 2010b). Social relationships in female chacma baboons (*Papio hamadryas ursinus*) help coping with stressful events and increase longevity (Silk et al., 2010a), whereas social bonds in male Assamese macaques (*Macaca assamensis*) and feral horses (*Equus ferus caballus*) are directly linked to the number of offspring sired (Schülke et al., 2010). However, it has also been observed that poor health could influence the social behaviour of group living animals through sickness-induced behavioural modulation (Adelman & Martin, 2009; Dantzer, O’Connor, Freund, Johnson, & Kelley, 2008; Klein, 2003)

In humans, a rich social network and social support yields measurable positive consequences for health (Berkman, 1995; Lisa. F. Berkman & Glass, 2000; Cassel, 1976; Cobb, 1976; Kiecolt-Glaser, McGuire, Robles, & Glaser, 2002; Theorell et al., 1995; Uchino, Cacioppo, & Kiecolt-Glaser, 1996). Recent research, however, has shown that complex sociality associated with the formation of strong, long-lasting social bonds, is not unique to humans and closely related primate species (Pollen et al., 2007; Shultz & Dunbar, 2010b). This suggests the fascinating hypothesis that positive health effects of social bonds may contribute to maintain stable group living in a large variety of taxa and types of social organizations (Silk, 2007).

Among birds, corvids show complex social relationships. Common ravens (*Corvus corax*), for example, are capable of forming valuable relationships, not only within reproductive pairs but also with social partners in their groups (Fraser & Bugnyar, 2012; Heinrich, 2011). Such bonds are characterized by low levels of aggression and high levels of affiliative behaviours exchanged in a reciprocal way over extended periods of time (Fraser & Bugnyar, 2010). Within such valuable relationships, individuals support each other in agonistic encounters (Emery, Seed, von Bayern, & Clayton, 2007; Fraser & Bugnyar, 2012) and share information and resources (Bugnyar, Kijne, & Kotrschal, 2001; de Kort, Emery, & Clayton, 2006; Fraser & Bugnyar, 2012), whereas they typically act competitively with other non-bonded conspecifics (Bugnyar & Heinrich, 2005, 2006). In the present study we investigated the benefits of social bonds in carrion crows (*Corvus corone*). In most European populations carrion crows form socially monogamous pairs during the breeding season (Glutz von Blotzheim, 1985; Meide, 1984) whereas in northern Spain crows live in stable social groups of up to nine individuals, consisting of the breeding pair and retained offspring as well as male immigrants (Baglione, Canestrari, Marcos, & Ekman, 2003). Group living in carrion crows is based on kinship (Baglione et al., 2003) and cooperation in nestling provisioning (Canestrari, Chiarati, Marcos, Ekman, & Baglione, 2008) and territory defence (Hillemann, et al. n.d.). Dominant males are more aggressive towards same sex group members compared to females (Chiarati, Canestrari, Vera, Marcos, & Baglione, 2010) and are less aggressive towards related individuals than to unrelated ones (Chiarati, Canestrari, Vila, Vera, & Baglione, 2011). In previous experiments, attentiveness of captive crows towards visual or olfactory social stimuli was measured. Males watched non-kin more often than kin individuals (Wascher, Núñez Cebrián, Valdez, Canestrari, & Baglione, 2014) and, unlike females, showed less avoidance towards the scent of a stressed familiar individual, compared to a stressed unfamiliar one, which might reflect a stronger willingness to provide social support (Wascher, Heiss, Baglione, & Canestrari, 2015).

In this study, we investigated the correlation between the quality of individual’s social relationships and the prevalence of gastrointestinal parasites in captive groups of carrion crows that varied in size and composition. In particular, we focused on the pattern of excretion of oocysts of coccidian protozoans and different nematode species, mainly *Syngamus trachea* and *Capillaria sp*. Coccidian oocysts are among the most common endoparasites of birds (López, Figuerola, & Soriguer, 2007; Page & Haddad, 1995) and are widespread among corvids, including carrion crows (Cawthorn & Wobeser, 1985; Poon & Chew, 1991; Upton, Langen, & Wright, 1995). Infections with large numbers of coccidia clinically manifest as ‘coccidiosis’, which damages the intestinal tract (Conway & McKenzie, 2007) and severely affects individual body condition, longevity and fecundity by inhibiting the uptake of essential dietary components (Hõrak et al., 2004; Stenkewitz, Nielsen, Skírnisson, & Stefánsson, 2016). The pathogenicity is well documented in poultry (Allen, 1987; Allen & Fetterer, 2002; Allen, Lydon, & Danforth, 1997), and further evidence is accumulating in wild bird species (*e.g*. Brawner, Hill, & Sundermann, 2000; McGraw & Hill, 2000). Coccidia are transmitted via the faecal-oral route, when oocysts undergo sporogony and become infectious (typically within 24 hours after excretion) (Allen & Fetterer, 2002). Similarly, different nematode species are described as regular endo-parasites of different corvid species (Halajian et al., 2011; Loman, 1980). *Syngamus trachea* is usually located in the trachea of birds and can lead to severe respiratory distress and pulmonary lesions (Fernando, Stockdale, & Remmler, 1971; Nevarez, Gamble, Aczm, Tully, & Avian, 2002). *Capillaria* sp. infect the intestinal tract and clinical symptoms are weight loss, diarrhoea, regurgitation, anaemia and oral necrotic plaques (Schnieder, Boch, & Supperer, 2006). Different nematode species infect their avian hosts via a direct life cycle, when embryonated eggs are ingested. Nematode eggs have a prepatent period of several weeks (Ortlepp, 1923).

In the present manuscript we ask how social factors relate to parasite egg and oocysts excretion patterns. First, we ask whether group size is related to parasite eggs and oocysts excretion, as infections may spread more easily in crowded conditions. Second, we ask whether parasite eggs and oocysts excretion correlate with a) the quality of the social relationships, with individuals maintaining strong social bonds being less likely to be infected than those with weak social bonds; b) the presence of relatives in the group, through reduction of stressful conflicts and competition (Chiarati et al., 2011) that can impair the ability to fight against parasites (Schat & Skinner, 2008); and c) an individual’s position in the dominance hierarchy, as dominance rank is known to weaken the immune system either in subordinates or dominants, depending on the relative allostatic load of social status in the respective species (Goymann & Wingfield, 2004).

## Methods

### Ethical Note

All procedures described in this manuscript were conducted in accordance with the guidelines for the treatment of animals in behavioural research of the Association for the Study of Animal Behaviour. Keeping crows in captivity and the procedures performed were authorized by Junta de Castilla y León (núcleo zoológico 005074/2008; EP/LE/358/2009).

### Study subjects

We collected data during seven different phases (November 2008, December 2008 - January 2009, January - February 2013, May - June 2013, January - February 2014, May – June 2014, September – October 2014), from a total of 36 crows (17 females, 19 males), which were housed in a large outdoor aviary (30 x 12 x 6 m) in Northern Spain. The aviary was located in Solanilla, León in 2008 and 2009 (42°37’25” N, 5°26’58” W), and moved location to Navafría, León in 2013 and 2014 (42°36’15” N, 5°26’56” W). The outdoor aviary resembled a natural situation as much as possible, with external walls and ceiling made of wire-net. It comprised four major compartments (12 x 6 x 3 m) separated by wooden walls and connected through eight smaller testing compartments (3 x 3 x 3 m). Compartments were equipped with wooden perches, natural vegetation, rocks, and sheltered perches in the four corners, where the wire-net ceiling was covered with transparent plastic boarding. There was no artificial floor, but natural vegetation covered the ground. An enriched diet consisting of fruit, vegetables, bread, meat and milk products was provided on a daily basis. Water and dry food were available *ad libitum*.

Birds were kept in groups that mirrored the social aggregations that naturally occur in the wild (A) ‘Flock’: three or more juvenile individuals; (B) ‘Pairs’: adult male and adult female. (C) ‘Cooperative Family’: a reproductive pair with its own already independent offspring. In 2008, 2010 and 2012 crows were hand-raised and initially kept in juvenile flocks. When birds aged and needed to be separated because of increasing frequency of aggressions, this was decided according to observed behaviour in the flock, *i.e*. individuals showing high frequency of affiliative behaviour and spatial proximity where paired. In June 2008, two wild families were captured in their territory at Sobarriba, brought into the aviary and subsequently kept in captivity (for more details, see: Wascher, Núñez Cebrián, Valdez, Canestrari, & Baglione, 2014). Each group of crows was kept in a different compartment, with acoustic but not visual contact with the rest of the captive birds (supplementary figure 1). Birds changed over time (from 2008 to 2014), so new groups were housed in separate compartments of the aviary. In each phase of data collection, each group was housed in only one keeping compartment of the aviaries. Between phases, group composition, size and holding compartment could change (supplementary materials Table 1). Although birds did not move freely between compartments and therefore were spatially isolated from each other, they were all equally potentially exposed to parasite transmission as it occurs in the wild, due to the outdoor settings and lack of hygienic isolation of each compartment. Droppings were removed from the aviary on a regular basis, approximately every couple of months, but no disinfection was applied.

### Behavioural data

A total of 1180 individual focal observations were recorded. Each observation lasted 5 minutes, and all occurring behaviours were recorded. For this study, we focused on frequencies of agonistic behaviour (threat, chase flight, and fight) and affiliative behaviours (allopreening and sitting in contact). The identity of the interacting individuals was noted, as well as their role (initiator/receiver) and the outcome of the interaction (winner/loser). The loser of an interaction was defined as the individual retracting. All behavioural observations were recorded on video and analysed by one researcher (C.A.F.W.).

### Social bonds: the composite sociality index

For each phase of data collection, we calculated a composite sociality index (CSI) for each crow dyad within a group according to Silk et al. (2010a). A CSI was constructed from two affiliative behaviours: contact sitting and allopreening. Note that we used mean values per observation instead of absolute numbers because the number of focal observations varied among individuals (supplementary Table 1). The higher the CSI of a dyad, as compared with the frequency of affiliative interactions observed in its entire group, the stronger the affiliative bond between the two individuals involved. Dyads with CSI higher than the average of the entire sample and lower rates of aggression compared to the average of the entire group, were classified as ‘bonded’. For each individual we calculated the maximum CSI value among all the dyads, reflecting the strongest affiliative relationship for each individual in the group.

### Agonistic encounters: Elo-rating

We calculated the success of individuals in agonistic encounters using Elo-rating in ‘aniDom’ (Sánchez-Tójar et al. 2017). Elo-rating allows to track dynamic changes in rank over different phases of data collection, by rating each of the individuals depending on the outcome of each single interaction (won or lost) and the probability of that outcome occurring (Neumann et al. 2011).

### Parasitological examination

During the entire study period, 760 individual droppings were collected on 160 days of sample collection. A human observer was watching the crows from inside the aviary and opportunistically collected droppings directly after defecation, so that each sample could be assigned to a particular individual (supplementary Table 1). To avoid diurnal variation in parasite eggs and oocysts shedding, we only collected droppings in the morning between 0900-1200. For each individual, a maximum of three droppings were collected on each sampling day (mean ± SE = 1.256 ± 0.238). Samples were stored in a refrigerator and analysed within seven days after collection. Samples were examined for eggs and oocysts of intestinal parasites. In 2008, we used a modified version of the flotation method (Schnieder et al., 2006) to examine the occurrence of parasite eggs and oocysts in the droppings. Fresh droppings (0.1 g) were suspended in a 2 ml collecting tube with 1 ml saturated saline. Collection tubes were shaken for 10 seconds and afterwards centrifuged for 5 minutes at 3000 rpm. After centrifugation, the collection tubes were filled with saline solution and a cover slip (18 x 18 mm) was positioned onto the tube. The high density of the saline solution causes the parasite eggs and oocysts to float up and be caught on the cover slip (Carta & Carta, 2000). After 10 minutes, the cover slip was moved onto a microscope slide and the parasite eggs and oocysts were identified and counted. From January 2013 onwards, we used a two grid McMaster (Marienfeld) counting chamber to examine the occurrence of parasite eggs and oocysts in the droppings. The entire sample was weighed, then diluted with 3 ml saturated NaCl solution per 0.1 g of droppings and thoroughly mixed. Afterwards, the solution was filled into both McMaster counting chambers. After 10 minutes of resting period we identified the parasite eggs and oocysts in both chambers. We used a compound microscope with 100-fold and 400-fold amplification to identify parasite eggs and oocysts. We found Coccidian oocysts, eggs of several nematode species (*Capillaria sp., Ascarida sp., Syngamus sp., Heterakis sp., Trichostrongylus tenius*) and cestode eggs to a varying degree. The proportion of positive samples was highest for coccidian oocysts (31 %, *N* = 760). Nematode eggs were found in 9 % of samples and cestode eggs where only found in less than 1 % of samples. We therefore limited our statistical analysis to coccidian oocysts and nematode eggs and analysed their occurrence (presence/absence) in the droppings, a measure that is unaffected by the sampling method used (Cringoli et al., 2011; Rinaldia, Coles, Maurelli, Musella, & Cringoli, 2011).

### Data analysis

We analysed the factors affecting the proportion of droppings containing coccidian oocysts and nematode eggs in our housed crows using the *glmer* function using R v. 3.0.2 (R Core Team 2016) and the glmer function in the *lm4* package (version 1.1-19; Bates, et al. 2014). We calculated a GLMM with binomial error distribution and a two-vectors response variable comprising the number of infected and non-infected samples for each individual in any given period of data collection. Various model diagnostics were employed to confirm model validity (visual inspection of distribution of residuals, qq plots, residuals plotted against fitted values) none of which suggested violation of model assumptions. To assess multicollinearity between fixed factors, we calculated variance inflation factors (VIFs) using the vif function in the package car (Fox & Weisberg, 2011). VIFs for all factors were below 1.5, indicating that there was no issue with multicollinearity (Zuur, Ieno, Walker, Saveliev, & Smith, 2009). We based our model selection on second-order Akaike’s Information Criterion values (AICc; Hurvich, Tsai, & Chih-Ling, 1989). We calculated the difference between the best model and each other possible model (ΔAICc) and ranked the model combinations according to their ΔAICc, which provides an evaluation of the overall strength of each model in the candidate set. Multiple models qualified as the similarly good models, *i.e*. ΔAICc ≤ 2 (Burnham, 2004; Burnham & Anderson, 2002) and therefore we applied a model averaging approach, which calculates model averaged parameters using the MuMIn package (version 1.15.6; Bartón, 2014). Full statistical models are presented in the supplementary materials (supplementary Table 2 and 3). Maximum CSI value, group size, number of related individuals, Elo-rating and sex were included as explanatory variables. For each model, we fitted individual identity as a random term to control for the potential dependence associated with multiple samples from the same individuals. To describe the variance explained by our models, we provide marginal and conditional R^2^ values that range from 0 to 1 and describe the proportion of variance explained by the fixed and by the fixed and random effects combined, respectively (Nakagawa & Schielzeth, 2013). We calculated marginal and conditional R^2^ values using the r.squaredGLMM function in MuMIn. Levels of significance were adjusted to *P* ≤ 0.025 according Bonferroni, to account for multiple testing of coccidia oocysts and nematode eggs.

## Results

### Social bonds

On average ± standard error (SE) we recorded 23.64 ± 3.38 affiliative interactions per individual as well as 12.21 ± 2.41 interactions won and 17.51 ± 2.44 lost per individual. We observed 56 bonded dyads (out of 327 dyads in total), 37 of which were male-female dyads (17 between related individuals and 20 between unrelated individuals), 11 were male-male (7 between related and 4 between unrelated individuals) and 8 female-female (7 related, 1 unrelated). Thirty-three dyads with CSI higher than the average of the entire sample but also higher rates of aggression, where not classed as bonded. On average ± SE, males and females had 1.58 ± 0.41 and 1.53 ± 0.36 bonded partners respectively. Mean CSI ± SE between bonded dyads was 2.73 ± 0.46 for male-female dyads, 4.06 ± 0.76 for female-female bonds and 3 ± 0.87 for male-male bonds.

### Occurrence of coccidian oocysts

Overall, 235 samples from 26 individuals contained coccidian oocysts, out of a total of 760 samples. Coccidian oocysts occurred in 151 out of 459 samples in males (33 %) and 84 out of 301 samples in females (28 %). Maximum CSI value, the number of related individuals in the group, Elo-rating score and sex remained as fixed factors in the best models (Table 1). Crows with a strong affiliative relationship (maximum CSI value: estimate ± SE = −0.11 ± 0.02, *z* = 4.28, *P* < 0.001) excreted a lower proportion of samples containing coccidian oocysts (Figures 1). A similar pattern occurred for individuals living in larger groups (estimate ± SE = −0.07 ± 0.01, *z* = 4.59, *P* < 0.001) and with more related individuals (estimate ± SE = −0.07 ± 0.03, *z* = 2.6, *P* = 0.01). On average ± standard error (SE) we recorded 12.21 ± 2.41 interactions won and 17.51 ± 2.44 lost per individual. Dominance rank (estimate ± SE = 0.06 ± 0.03, *z* = 1.81, *P* = 0.07) and sex (estimate ± SE = −0.01 ± 0.04, *z* = 0.33, *P* = 0.74) did not significantly relate to the proportion of samples containing coccidian oocysts.

**Table 1.** Model selection of analyses that examined the occurrence of coccidian oocysts, related to maximum CSI value (max CSI), number of related individuals (nr related), Elo-rating scores, group size and sex.

| | Models | df | LogLik | AICc | ΔAICc | Weight | $R^2_{\text{marginal}}$ |
| --- | --- | --- | --- | --- | --- | --- | --- |
| (a)<br>*Coccidia* | Max CSI, Elo-rating, group size | 5 | -165.45 | 341.7 | 0.00 | 0.15 | 0.0 |
|  | Max CSI, Elo-rating, nr related | 5 | -165.58 | 341.95 | 0.25 | 0.13 | 0.09 |
|  | Max CSI, Elo-rating, group size, nr related | 6 | -164.44 | 342 | 0.31 | 0.13 | 0.09 |
|  | Max CSI, group size | 4 | -167.07 | 342.66 | 0.96 | 0.09 | 0.06 |
|  | Max CSI, group size, nr related | 5 | -166.14 | 343.07 | 1.37 | 0.08 | 0.08 |
|  | Max CSI, nr related | 4 | -167.32 | 343.16 | 1.46 | 0.07 | 0.07 |
|  | Max CSI, Elo-rating, group size, sex | 6 | -165.26 | 343.64 | 1.94 | 0.06 | 0.06 |
| (b)<br>*Nematodes* | Null model | 2 | -103.42 | 210.99 | 0 | 0.16 | 0 |
|  | Sex | 3 | -102.57 | 211.46 | 0.47 | 0.13 | 0.02 |
|  | Elo-rating | 3 | -103.27 | 212.85 | 1.86 | 0.06 | 0 |

Individual identity was fitted as a random term in all models. LogLik: log-likelihood, df: degrees of freedom, ΔAICc: difference in AICc to the best model, marginal and conditional $R^2$ . Models are ranked according to their Akaike weight (Weight).

**Table 2.**
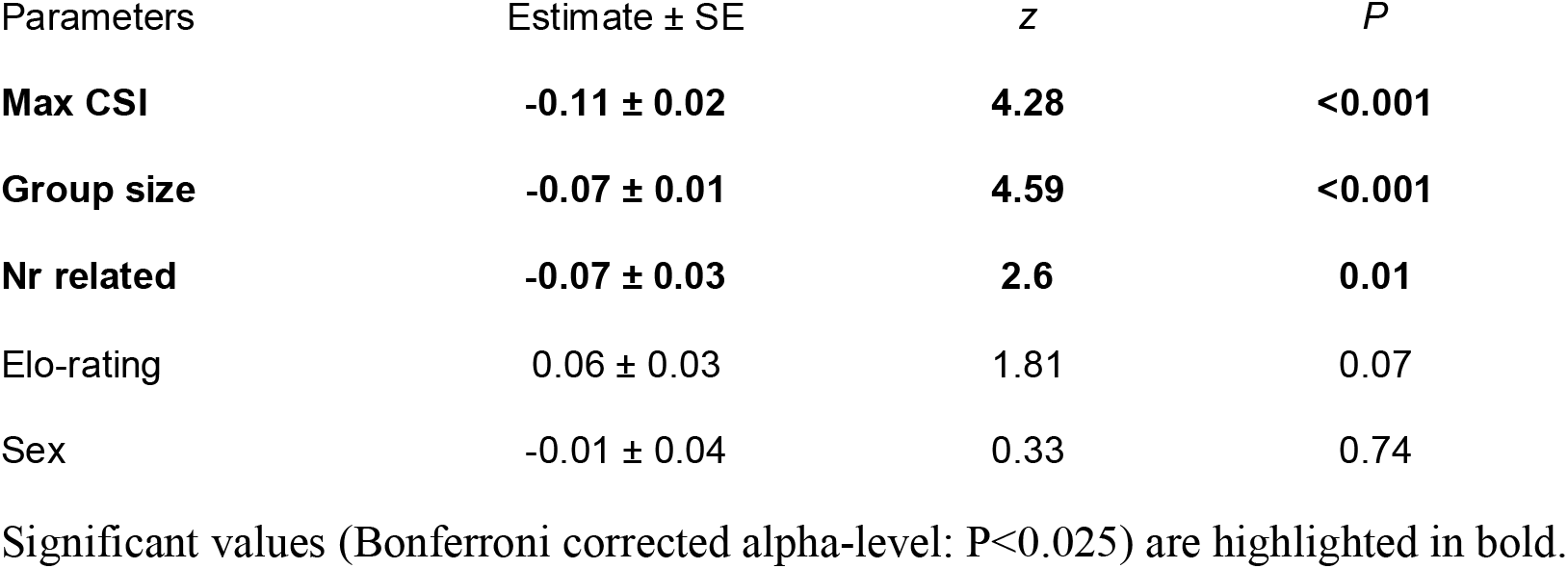
Results of model averaged generalized linear mixed model investigating factors affecting patterns of coccidian oocyst excretion.

**Figure 1.**
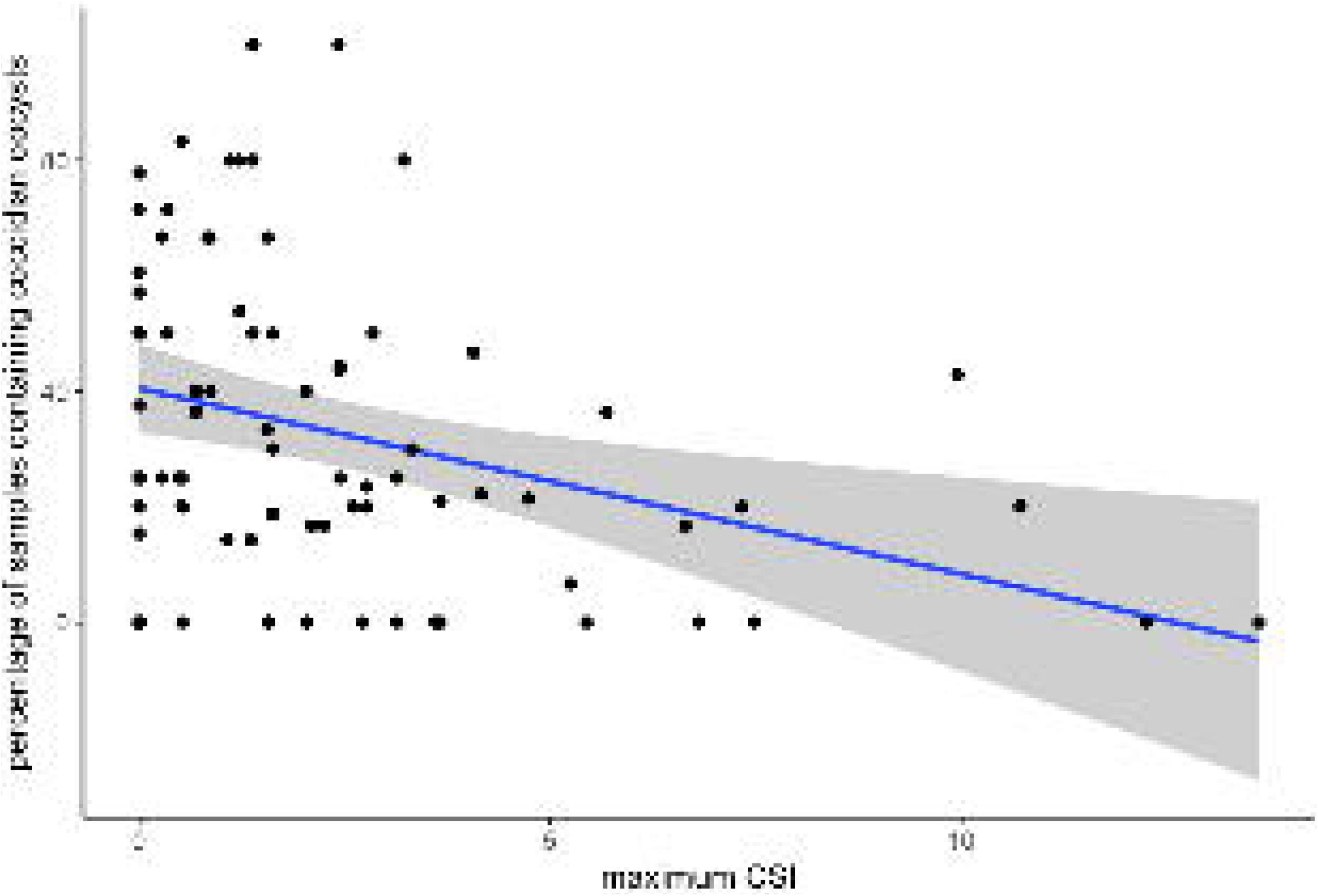
Percentage of samples containing coccidan oocysts in carrion crow droppings in relation to the maximum CSI. The predicted values are shown as solid line and 95% CI as shaded area. Black dots present individual values per phase.

### Occurrence of nematode eggs

Overall, 69 samples from 23 individuals contained nematode eggs, out of the total of 760 samples collected in all the 36 individuals. None of the factors investigated significantly affected excretion patterns of nematode eggs (supplementary Table 3), in fact the null model was amongst the best models in the candidate set (Table 1).

## Discussion

In the present study, we have shown that carrion crows with closely bonded social partners (high CSI scores) were less likely to excrete coccidian oocysts, which represent an important threat for health in birds (Hõrak et al., 2004; Stenkewitz et al., 2016). Health correlates of high quality social ties have been reported in humans (e.g. Seeman, 1996). Similar evidence has been recently found in non-human primates (Schülke et al., 2010; Silk et al., 2010a), suggesting that the tendency to form strong social ties may have deep phylogenetic roots (Shultz & Dunbar, 2010a). Social complexity, however, is not limited to primate species. Indeed, coalition formation has been reported in several mammalian and bird species, e.g. African elephants (*Loxodonta africanus*) (Bates et al., 2008), spotted hyenas (*Crocutta crocutta*) (Holekamp, Sakai, & Lundrigan, 2007), bottlenose dolphins (*Tursiops aduncus*) (Connor & Krützen, 2015), ravens (Braun & Bugnyar, 2012), and carrion crows (Baglione et al., 2003). Our study uncovered that high quality social relationships correlate with reduced occurrence of parasites. From our present observational study, we cannot conclude about the causality of effects. Parasite burden could either be affected by social bonds showing a benefit of social relationships in group living animals, or could influence the social behaviour through sickness-induced behavioural modulation (Adelman & Martin, 2009; Dantzer, O’Connor, Freund, Johnson, & Kelley, 2008; Klein, 2003), or sickness-induced cognitive biases (Nettle & Bateson, 2012). Either way, our data illustrate the importance of physiological mechanisms underlying social behaviours and potential benefits associated to the social environment in birds that parallel those of primates (including humans) and suggesting a route towards advanced sociality that may be common to a variety of taxa.

Cooperatively breeding groups of carrion crows form through two different mechanisms, namely delayed dispersal of offspring, which remain in the natal territory with their parents and siblings (Baglione, Marcos, & Canestrari, 2002), and formation of long lasting social bonds among distant relatives, most often males, that share all-purpose territories and frequently mate polyandrously (Baglione et al., 2003; Baglione, Marcos, Canestrari, & Ekman, 2002). These bonds typically form when an immigrant male joins an established family group and allies with the same sex resident breeder, to whom it is related (Baglione et al., 2003). Cooperative breeding, with group members working together to raise the brood, can only arise once stable groups have formed (Canestrari, Marcos, & Baglione, 2005). Both offspring delayed dispersal and bonding between adult males are therefore necessary preceding steps that eventually lead to cooperation, which has to be considered a consequence rather than the cause for the formation of the social group (Ekman, Baglione, Egger, & Griesser, 2001; Ekman, Dickinson, Hatchwell, & Griesser, 2004; Hatchwell & Komdeur, 2000). In other words, to understand why carrion crows, as well as any other bird species, form groups we need to understand the advantages that sociality conveys, independently of the payoff derived from cooperation at the nest. In carrion crows, as well as in many other cooperatively breeding species, the benefits of delayed dispersal for offspring are well studied (Chiarati, Canestrari, Vera, & Baglione, 2012), but little is known about the advantages of forming long lasting social bonds outside the nuclear family. The correlation between social bonds and parasite burden found in this study is consistent with the idea that the health benefits of high-quality relationships extend beyond the bonds between parents and offspring or reproductive partners, and may be associated with a reduction of harmful endo-parasites. Ultimately, this may be an important factor for establishing stable relationships in groups. Further research is needed to confirm the direction of the cause and effect relationship of these results. However, it should be noted that besides an effect of strong social bonds, we also found that the number of relatives in the group was negatively correlated with parasite oocyst excretion in carrion crows. Because group size and composition in this captive situation were obviously not under the control of the crows themselves, the reduced parasite burden in kin-based groups was likely to be a consequence, and not a cause, of the presence of relatives, suggesting that social bonds in general affect health and not vice-versa. Sociality is based on kinship in wild cooperatively breeding carrion crows. Offspring remain in the natal territory with their parents for years and more distant relatives are actively recruited to form cooperative alliances with the resident breeders (Baglione et al., 2003). As a result, social groups in cooperatively breeding carrion crow are extended families, comprising members with different degree of relatedness. Indirect fitness benefits are known to be a primary driver of kin-based sociality in many taxa of animals (Clutton-Brock, 2002). However, our results indicate that living with kin can also accrue immediate direct benefits to carrion crows through reduced infection by coccidia.

Increased exposure to parasites and disease transmission is considered as one of the major disadvantages of group living (Côté & Poulin, 1995). In our study, however, group size showed the opposite effect, with the probability of presence of coccidian oocysts decreasing in larger groups. This suggests that the benefits of sociality in crows are not dumped by the health risk of living in group, at least in the range of group sizes tested in this study. However, crows can also aggregate in larger flocks, typically in winter, when they roost and forage communally (Sonerund et al., 2002). The trade-off between the benefits of sociality and risk of infection in these particular circumstances are yet to be assessed.

We only found effects of social factors in coccidia, but not in nematode species. One reason could be that the lower occurrence of nematode eggs compared to coccidian oocysts in the samples (12 % of samples contained nematode eggs and 24 % of samples contained coccidian oocysts) limited the power of our analysis. However, the different life cycles of different parasite species could also play a role. Coccidia sp. have a prepatent period of approximately one week, in contrast nematode species, *e.g*. capillaria, have a pre-patent period of 3 - 4 weeks (Schnieder et al., 2006). Therefore, we suggest that short-term changes in the immune system, which could be caused by suppressive effects of glucocorticoids in response to social interactions (Bartolomucci, 2007), might have stronger effects on infection by coccidia compared to nematode species.

The present study was conducted in captive, mostly hand-raised individuals, because this allowed for repeated observations of individuals habituated to the presence of a human observer. Working in aviaries is a standard procedure to investigate social behaviour in corvids (Fraser & Bugnyar, 2012; Kondo & Hiraiwa-Hasegawa, 2015; Logan, Ostojić, & Clayton, 2013; Wascher, 2015) and although similar observational studies in the wild are desirable, they are difficult to realize, because corvids avert humans and are difficult to observe in the wild. However, differentiated affiliative social relationships have previously been described in wild ravens (Braun & Bugnyar, 2012; Braun, Walsdorff, & Bugnyar, 2012) as well as carrion crows (Baglione et al., 2003) and types of social aggregations in the aviary, *e.g*. flock, pairs, families, reflect actual forms of social aggregations observed in the wild (Baglione, Marcos, & Canestrari, 2002). The investigated parasite species are widespread among wild corvids, including carrion crows (Cawthorn & Wobeser, 1985; Poon & Chew, 1991; Upton et al., 1995) and have been found in wild carrion crows at the study site in northern Spain (Wascher, 2014).

### Conclusions

Social complexity has deep phylogenetic roots, being widespread in a wide variety of mammals and birds. Understanding its evolution, however, is difficult because of our scant knowledge of the fitness consequences of sophisticated social behaviour across taxa. We are just starting to fill the gaps, and this study shows a correlative relationship between the quality of social relationships and parasite shedding in a bird species that may underlie social bonding, setting the stage for a complex form of cooperation (cooperative breeding).

## Acknowledgments

We thank Kurt Kotrschal for feedback and discussion, Hugo Robles for statistical advice and Friederike Hillemann for help with droppings collection. Alfredo Sánchez-Tójar advised on calculation of dated individual Elo-rating trajectories. Further we would like to thank Bernhard Voelkl and two anonymous reviewers for valuable comments on earlier versions of the manuscript. This work was supported by the Spanish Plan Nacional I+D, project CGL2011-27260 and CGL2016-77636-P, to VB and a KWA (‘kurzfristiges wissenschaftliches Auslandsstipendium’) stipend of the University of Vienna to C.A.F.W..

## Appendix

**Appendix Table 1:**
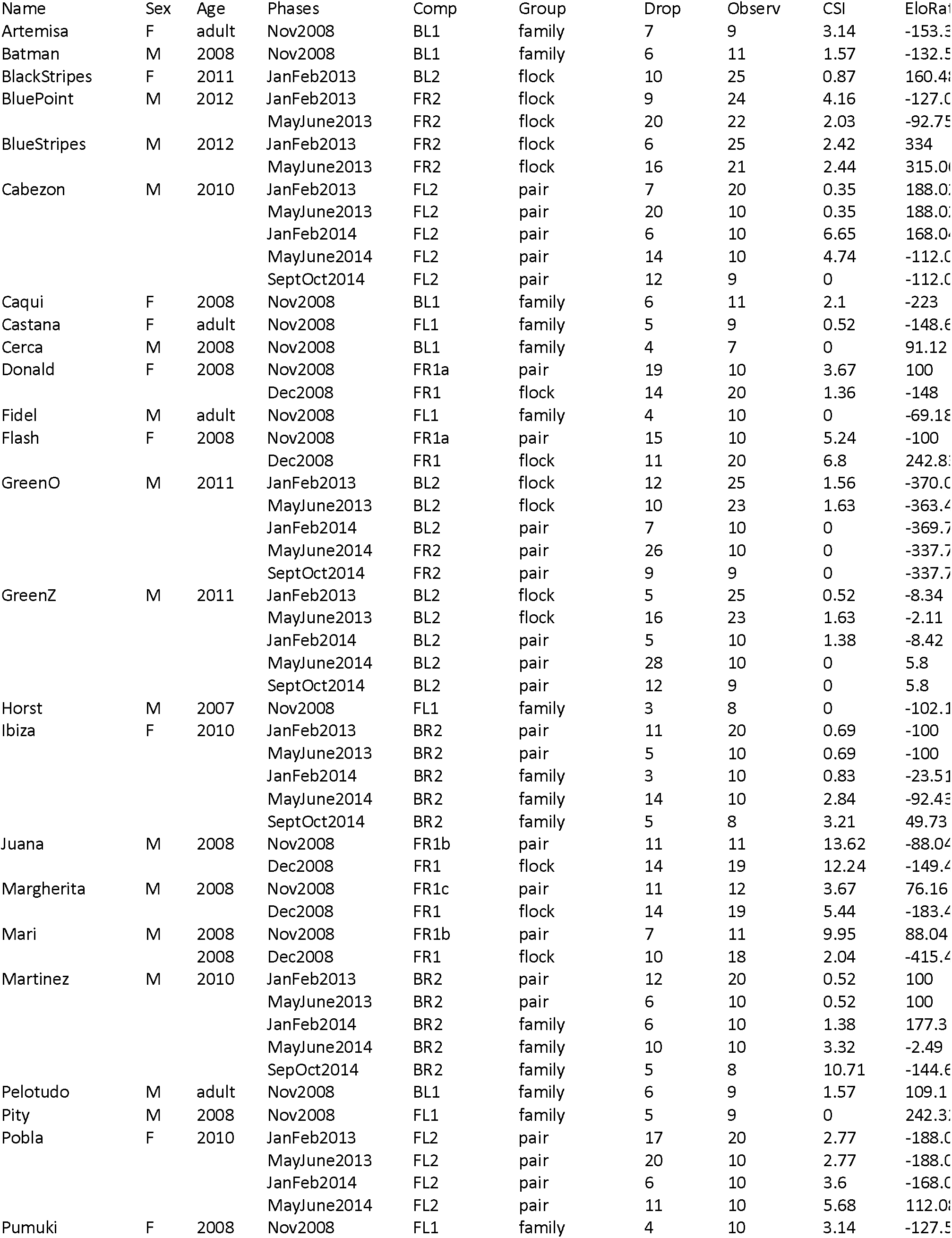
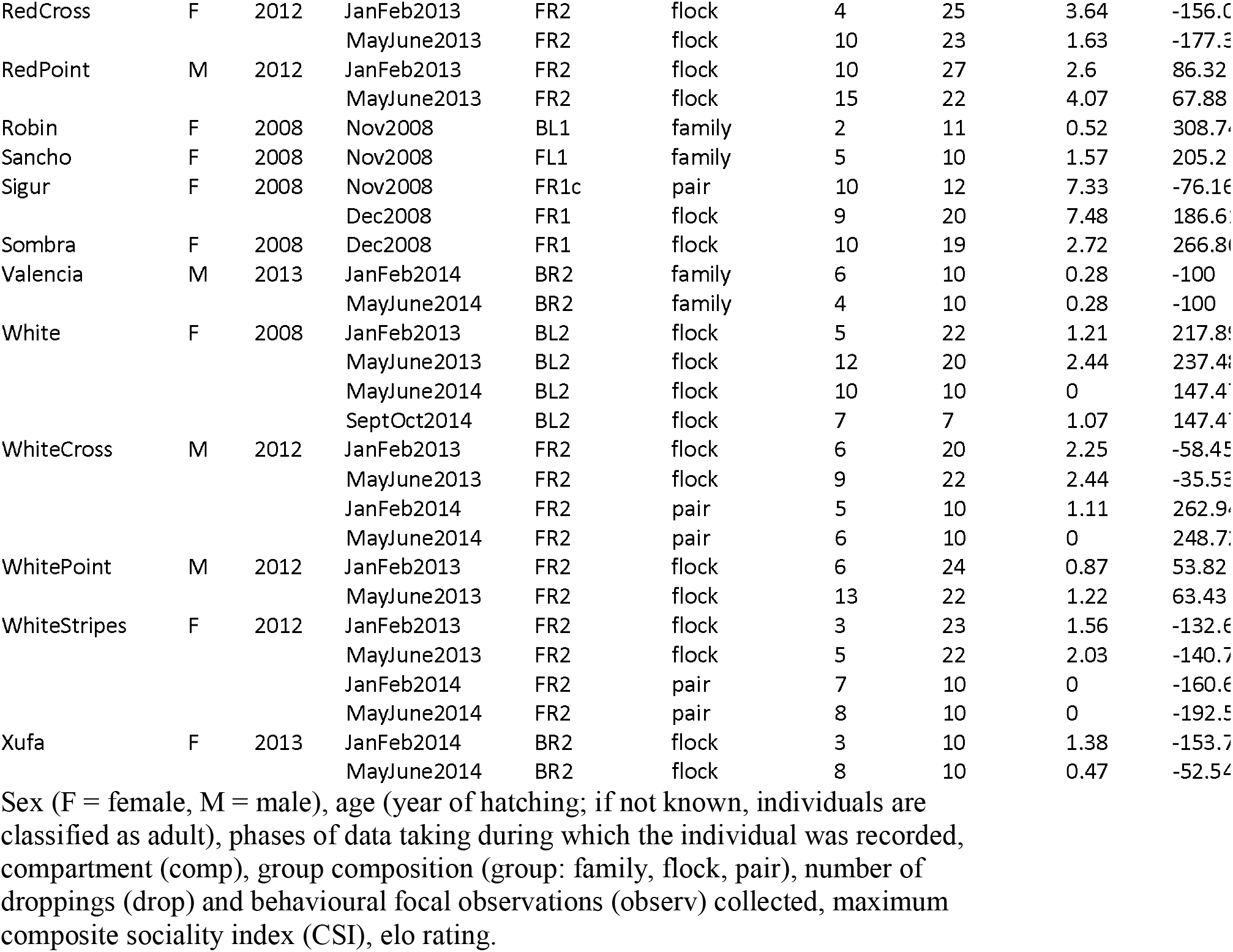
List of all focal individuals and background information

**Appendix Table 2:** Results of the full generalized mixed linear model investigating factors relating to patterns of coccidian oocyst excretion. Bonferroni adjusted significant values (*P* ≤ 0.025) are highlighted in bold.

| Parameters | Estimate ± SE | *z* | *P* | Odds<br>Ratio | Lower 2.5%<br>CI | Upper<br>97.5% CI |
| --- | --- | --- | --- | --- | --- | --- |
| Intercept | 0.1 ± 0.37 | 0.28 | 0.78 | -0.031 | -0.67 | 0.83 |
| **Maximum CSI** | **-0.15 ± 0.05** | **-2.84** | **0.004** | **-0.14** | **-0.26** | **-0.04** |
| Group Size | -0.11 ± 0.07 | -1.5 | 0.13 | -0.03 | -0.26 | 0.03 |
| Number of<br>related<br>individuals | -0.15 ± 0.1 | -1.48 | 0.14 | -0.17 | -0.36 | 0.06 |
| Sex | -0.2 ± 0.32 | -0.64 | 0.52 | -0.14 | -0.88 | 0.46 |
| Elo-rating | 0 ± 0 | 1.85 | 0.06 | 0 | -0 | 0 |

**Appendix Table 3:** Results of the full generalized mixed linear model investigating factors relating to patterns of nematode egg excretion. Bonferroni adjusted significant values (*P* ≤ 0.025) are highlighted in bold.

| Parameters | Estimate ± SE | *z* | *P* | Odds Ratio | Lower 2.5% CI | Upper 97.5% CI |
| --- | --- | --- | --- | --- | --- | --- |
| **Intercept** | **-3.19 ± 0.54** | **-5.96** | **<0.001** | **-3.19** | **-4.27** | **-2.13** |
| Maximum CSI | 0.02 ± 0.07 | 0.34 | 0.72 | 0.02 | -0.13 | 0.16 |
| Group Size | 0.03 ± 0.08 | 0.44 | 0.66 | 0.03 | -0.12 | 0.19 |
| Number of related individuals | -0.02 ± 0.15 | -0.14 | 0.89 | -0.02 | -0.35 | 0.27 |
| Sex | 0.68 ± 0.48 | 1.41 | 0.16 | 0.68 | -0.35 | 1.62 |
| Elo-rating | -0 ± 0 | -0.58 | 0.56 | -0 | -0 | 0 |

**Figure A1.**
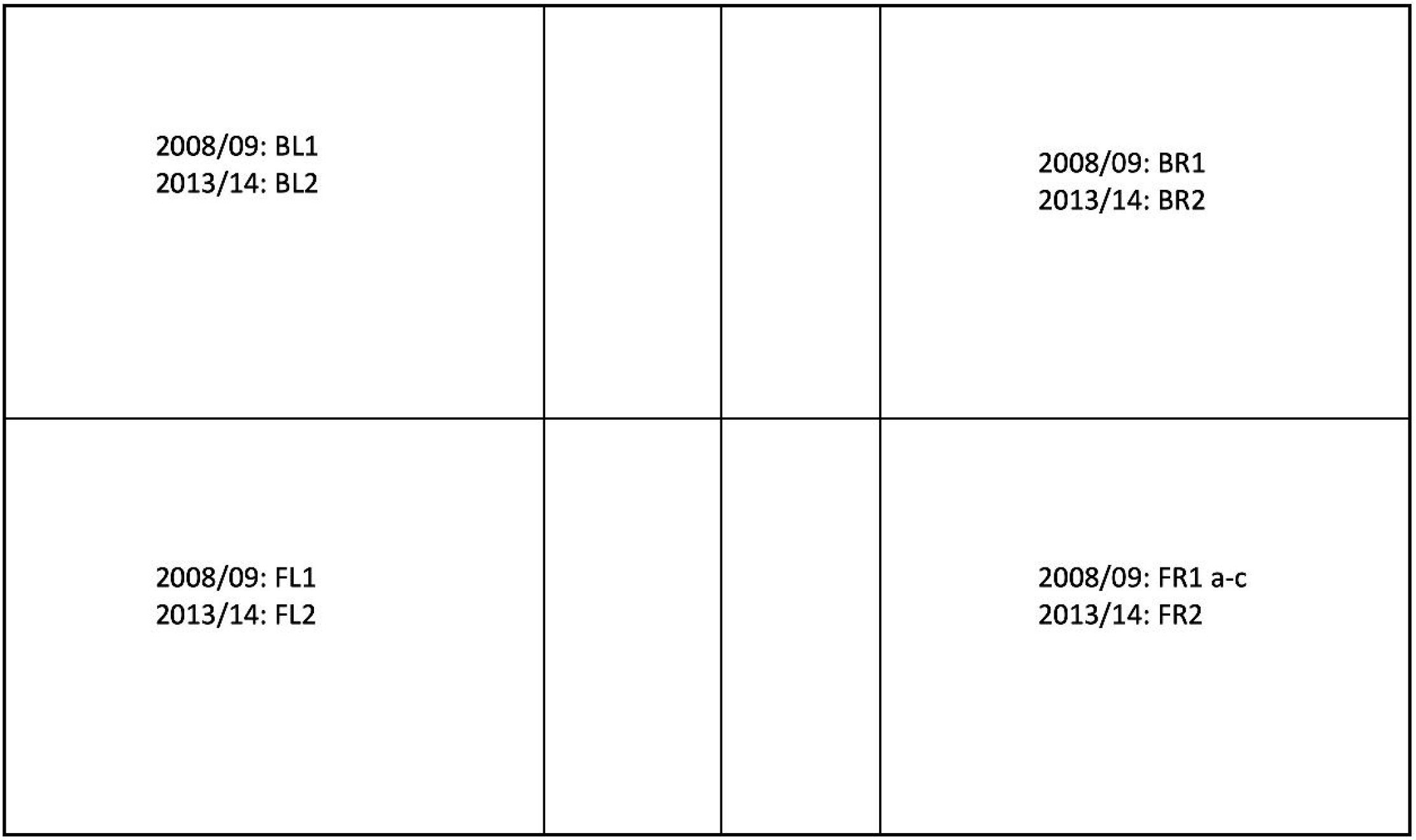
Sketch of the aviary setup. Aviary consisted of four separate compartments, in visual, but not acoustic separation from each other. Aviary was kept in two separate locations, in La Solanilla, León in 2008 and 2009 (42°37’23.4336 N, 5°27’3.1788 W), and Navafría, León in 2013 and 2014 (42°36’33, N 5°26’56 W).

